# Multiple TonB Homologs are Important for Carbohydrate Utilization by *Bacteroides thetaiotaomicron*

**DOI:** 10.1101/2023.07.07.548152

**Authors:** Rebecca M Pollet, Matthew H Foley, Supriya Suresh Kumar, Amanda Elmore, Nisrine T Jabara, Sameeksha Venkatesh, Gabriel Vasconcelos Pereira, Eric C Martens, Nicole M Koropatkin

## Abstract

The human gut microbiota is able to degrade otherwise undigestible polysaccharides, largely through the activity of the *Bacteroides*. Uptake of polysaccharides into *Bacteroides* is controlled by TonB-dependent transporters (TBDT) whose transport is energized by an inner membrane complex composed of the proteins TonB, ExbB, and ExbD. *Bacteroides thetaiotaomicron* (*B. theta*) encodes 11 TonB homologs which are predicted to be able to contact TBDTs to facilitate transport. However, it is not clear which TonBs are important for polysaccharide uptake. Using strains in which each of the 11 predicted *tonB* genes are deleted, we show that TonB4 (BT2059) is important but not essential for proper growth on starch. In the absence of TonB4, we observed an increase in abundance of TonB6 (BT2762) in the membrane of *B. theta*, suggesting functional redundancy of these TonB proteins. Growth of the single deletion strains on pectin galactan, chondroitin sulfate, arabinan, and levan suggests a similar functional redundancy of the TonB proteins. A search for highly homologous proteins across other *Bacteroides* species and recent work in *B. fragilis* suggests that TonB4 is widely conserved and may play a common role in polysaccharide uptake. However, proteins similar to TonB6 are found only in *B. theta* and closely related species suggesting that the functional redundancy of TonB4 and TonB6 may be limited across the *Bacteroides*. This study extends our understanding of the protein network required for polysaccharide utilization in *B. theta* and highlights differences in TonB complexes across *Bacteroides* species.

**Importance:** The human gut microbiota, including the Bacteroides, is required for the degradation of otherwise undigestible polysaccharides. The gut microbiota uses polysaccharides as an energy source and the fermentation products such as short chain fatty acids are beneficial to the human host. This use of polysaccharides is dependent on the proper pairing of a TonB protein with polysaccharide-specific TonB-dependent transporters; however, formation of these protein complexes is poorly understood. In this study, we examine the role of 11 predicted TonB homologs in polysaccharide uptake. We show that two proteins, TonB4 and TonB6, may be functionally redundant. This may allow for development of drugs targeting *Bacteroides* species containing only a TonB4 homolog with limited impact on species encoding the redundant TonB6.

## Introduction

The human gut microbiota performs many important functions that promote human health including the degradation of complex carbohydrates (fiber or polysaccharides) from our diet (1). Many bacteria in the microbiota ferment polysaccharides, resulting in the release of short-chain fatty acids (SCFAs) such as butyrate, acetate, and propionate (2). These SCFAs then serve as a key energy source for colonocytes and promote intestinal barrier function (3, 4). Understanding the molecular mechanisms of polysaccharide degradation will provide opportunities to develop functional foods as therapeutics or inhibitors of polysaccharide degradation to manipulate microbial metabolism and improve human health.

The Gram-negative Bacteroidetes are abundant members of the Western adult gut microbiota and maintain a large capacity to degrade polysaccharides (5, 6). In these bacteria the genes required for polysaccharide use are organized into polysaccharide utilization loci (PUL) (7). Each PUL is generally transcriptionally activated in response to a distinct polysaccharide substrate (8, 9). Several lipoproteins encoded within each PUL localize to the cell surface to bind, degrade, and import the target substrate (7, 10). The prototypical PUL is the starch utilization system (Sus) from *Bacteroides thetaiotaomicron* (*B. theta*) (7, 11, 12). A common feature across all Sus-like systems is at least one pair of proteins homologous to SusC, a putative TonB-dependent transporter (TBDT), and SusD, a starch-binding protein (Fig 1) (7, 13). Detailed biochemical studies of the additional outer membrane proteins from PUL that target starch, arabinan, levan, chondroitin sulfate, heparin, and several other polysaccharides have helped develop a model of how polysaccharides are initially degraded at the cell surface and elucidate which oligosaccharides are selected and imported into the cell via the SusCD-like complex (14–21).

**Figure 1.**
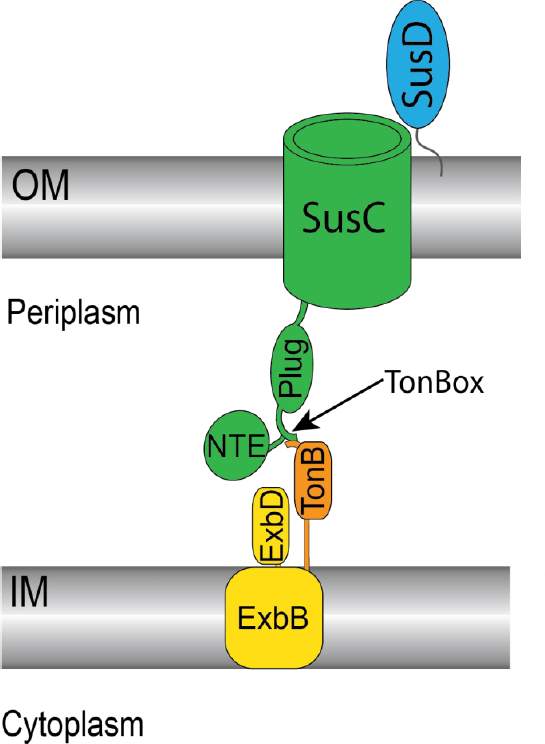
Architecture of the predicted 5usC, 5usD, TonB, and ExbBD complex. 5urface glycan binding protein 5usD is shown in blue, associated with 5usC at the outer membrane (OM}. TonB-dependent transporter 5usC is shown in green including the plug and N-terminal extension (NTE) domains. The TonBox precedes the plug domain and here is shown pairing with TonB (orange}. ExbB and ExbD are shown in yellow in the inner membrane (IM}.

SusC is required for starch utilization via transport of maltooligosaccharides across the outer membrane (22). SusC and its thousands of homologs in Bacteroidetes share sequence homology with well-studied TBDTs of Gram-negative organisms. Thus far, one TBDT from *Porphyromonas gingivalis* and three SusC-like transporters from *B. theta* have been structurally characterized and show high structural homology with TBDTs such as the well-studied FhuA and FepA from *Escherichia coli* (21, 23–25).

Key conserved features of these transporters include a 22 beta-strand barrel that traverses the outer membrane and houses a plug domain that occludes solute passage until the transporter is activated. The TonB box or TonBox is a sequence of 4-8 conserved residues forming a β-strand that precedes the plug domain and is important for pairing with TonB (Fig 1) (26). Some *B. theta* TBDTs are predicted to include two additional domains termed the N-terminal extension (NTE) and the Secretin and TonB N-terminus (STN) domain (25, 27). The precise rearrangement of the plug domain to allow solute passage through the barrel is poorly understood but is facilitated by pairing to an inner membrane complex that harnesses proton motive force. This complex includes the proteins TonB, ExbB, and ExbD (Fig 1). Structural analysis of this complex in *E. coli* and *Serratia marcescens* reveals the inner membrane spanning ExbB in a pentameric arrangement enclosed around a dimer of ExbD that extends into the periplasm (28, 29). The structure of the complex with TonB has not been determined but other characterization of TonB suggests at least one copy of TonB interacts with the ExbBD complex via the TonB N-terminal membrane spanning α-helix (30–32). This N-terminal α-helix is followed by a linker region that spans the periplasm and a well-ordered C-terminal domain (33). The C-terminal domains of characterized TonB proteins share a common fold of three antiparallel β-sheets with two α-helices (Fig 2BC) (34, 35). The final β-strand at the C-terminus directly contacts the TBDT for β-sheet pairing with the TonBox (Fig 2BC) (26, 36, 37).

**Figure 2.**
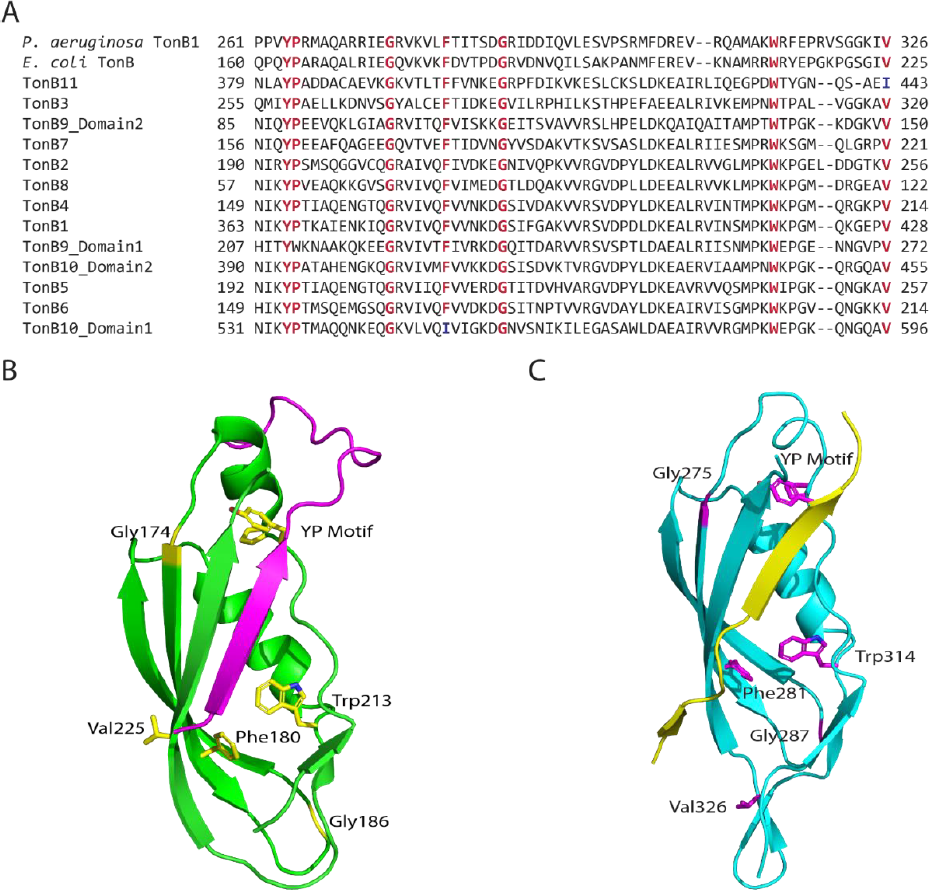
**A.** Excerpt of the multiple sequence alignment of the eleven identified TonB proteins from *B. thetaiotaomicron* with *E. coli* TonB and *Pseudomonas aeruginosa* TonB1. Conserved residues are highlighted in red, deviations from this conservation are highlighted in blue. The full alignment is shown in Fig. 52. **B.** 5tructure of the *E. coli* TonB (green) in complex with the TonBox of BtuB (magenta}, PDB: 2G5K Conserved residues from A are highlighted in yellow. **C.** 5tructure of *Pseudomonas aeruginosa* TonB1 (cyan) in complex with the TonBox of FoxA (yellow}, PDB: 6I97. Conserved residues from A are highlighted in magenta.

Evidence that transport through Bacteroidetes SusC-like systems is TonB dependent is supported by previous work in which the NanO sialic acid transporters from *Tannerella forsythia* and *Bacteroides fragilis* were functional in *E. coli* only in the presence of TonB (38). The xylan-targeting SusC-like protein from *Bacteroides vulgatus* was also shown to be functional in *E. coli* but dependence on TonB was not explored (39). Efforts to express SusC and other homologous *B. theta* transporters in *E. coli* have not been successful, preventing similar characterization (N.M. Koropatkin unpublished data, 13). Most recently, it has been shown that growth of *B. fragilis* on substrates known to be transported through TBDTs including heme, vitamin B12, iron, starch, mucin-glycans, and N-linked glycans is disrupted by deletion of a single *tonB* gene (40).

Unlike *E. coli* which expresses only one TonB, *B. theta* and other Bacteroidetes can encode up to 15 TonB homologs. In some Gram-negative organisms that encode multiple TonB proteins such as *Xanthomonas campestris*, there is evidence of TonB-TBDT pairing redundancy such that more than one TonB can energize a discrete transporter (41). Conversely, in *Caulobacter crescentus*, the deletion of a single TonB completely abrogates the import of maltose (42). TonB-TBDT pairing has been explored in two Bacteroidetes, *Riemerella anatipestifer* and *B. fragilis*. Characterization of the three TonB homologs in *R. anatipestifer* suggests each TonB functions differently but that there may be some redundancy with both TonB1 and TonB2 facilitating hemin uptake but loss of TonB2 having a much greater impact (43). Conversely, in *B. fragilis*, deletion of a single *tonB* gene completely eliminated growth on a variety of substrates including several different polysaccharides while deletion of the other 5 homologs had no impact on growth suggesting that a single TonB is responsible for pairing with a variety of transporters under these conditions (40).

The TonB proteins encoded by *B. theta* differ in both number and sequence from those encoded by *B. fragilis*, suggesting that TonB pairing may differ even between these closely related species. Through genetic analysis we identify 11 TonB homologs in *B. theta* and construct strains containing a gene deletion of each homolog. To better understand Sus as the prototypical PUL, we analyzed growth of each of these strains on starch and show that deletion of TonB4 leads to less efficient growth on starch. However, in contrast to *B. fragilis*, this deletion does not completely eliminate growth on starch or other polysaccharides, allowing us to identify a second, redundant TonB important in starch utilization. We then expand our analysis to other *B. theta* PUL with well-characterized TBDT showing a similar redundancy in TonB function. This work underscores the roles of TonB homologs during outer membrane transport and expands our understanding of glycan uptake in the *Bacteroides*.

## Results

### Identification of 11 TonB proteins in *B. theta*

High sequence diversity has been seen across characterized TonB proteins making high confidence annotation of TonB proteins difficult. In fact, varying number of genes are annotated as *tonB* across different analyses of the *B. theta* genome (13, 40, 44). We compiled a list of eleven potential TonB proteins in *B. theta* by searching the genome for proteins containing the Gram-negative bacterial TonB protein C-terminal Pfam domain (TonB_C, PF03544) (Table 1, additional information in **Fig S1A**). One additional protein (BT3921) showed a match to PF03544 but has not been included in this analysis due to the low confidence of that prediction (e-value = 2.7e-05). This analysis matches the eleven *B. theta* TonB proteins identified by Parker *et al.* (40).

**Table 1.** Identified TonB proteins in *Bacteroides thetaiotaomicron* VPI-5482.

| Protein Name | Locus Tag | Total Protein Length | Location of TonB_C Domain | E-value | Additional domain? | Location of additional domain | E-value of additional domain |
| --- | --- | --- | --- | --- | --- | --- | --- |
| TonB1 | BT0813 | 443 | 363-439 | 1.8e-21 | Peptidase_M56 (PF05569) | 137-266 | 5.7e-16 |
| TonB2 | BT1056 | 269 | 190-268 | 1.4e-14 | no |  |  |
| TonB3 | BT1668 | 350 | 256-331 | 4.5e-14 | no |  |  |
| TonB4 | BT2059 | 227 | 149-226 | 2.2e-23 | no |  |  |
| TonB5 | BT2665 | 270 | 192-269 | 1.6e-22 | no |  |  |
| TonB6 | BT2762 | 227 | 149-226 | 4.6e-23 | no |  |  |
| TonB7 | BT3192 | 249 | 156-232 | 1.0e-20 | CarbopepD-reg_2 (PF13715) | 33-112 | 2.5e-11 |
| TonB8 | BT3673 | 283 | 57-134 | 1.1e-20 | DUF4488 (PF14869) | 159-281 | 1.2e-40 |
| TonB9 | BT3896 | 285 | 207-284* | 1.3e-21 | TonB_C (PF03544) | 85-161 | 4.6e-21 |
| TonB10 | BT3898 | 609 | 531-608* | 1.1e-22 | TonB_C (PF03544) | 390-467 | 1.8e-18 |
|  |  |  |  |  | Peptidase_M56 (PF05569) | 160-261 | 3.2e-13 |
| TonB11 | BT4460 | 449 | 379-440 | 1.6e-07 | CarbopepD-reg_2 (PF13715) | 228-309 | 8.0e-19 |

\* For TonB9 and 10 which have two TonB domains, the domain with the lower E-value is used as the primary TonB domain. Domains are numbered beginning with the N-terminal most domain for subsequent analysis so these primary TonB domains are annotated as the TonB\_C2 domain in Fig. S1.

Sequence similarity between these eleven *B. theta* TonB proteins and *E. coli* TonB is low, ranging from 10-37% sequence identity (**Fig S1B**). This is partially due to lack of the TonB polyproline region (PF16031) in all candidate *B. theta* TonB proteins. In *E. coli* TonB, this region has been shown to be important, though not essential, for properly spanning the periplasm to interact both with the ExbBD complex and TBDTs (33, 45). Characterized TonB proteins with polyproline regions are predominantly from Enterobacteriaceae so the lack of this region in our identified TonB proteins may suggest a different structure is needed to properly span the periplasm of other bacteria including those from the Bacteroidetes phylum (33, 46, 47).

Several *B. theta* TonB proteins (TonB1, 7, 8, 9, 10, and 11) also contain additional domains appended to the TonB C-terminal domain (Table 1). To our knowledge TonB proteins with additional domains have not been functionally characterized so it is difficult to predict if these additional domains confer additional or altered functionality. The predicted peptidase domains (PF05569) in TonB1 and 10 could suggest a role for these proteins in signal transduction. The peptidase M56 family includes the BlaR1 protein that serves as the sensor-transducer for the β-lactam antibiotic resistance pathway in Staphylococci; however, there is no evidence of a similar mechanism for β-lactam resistance in *B. theta* (48, 49). Previous bioinformatic analysis of TonB proteins identified only nine proteins with this M56 N-terminal extension out of the 263 sequences analyzed (44). These nine TonB proteins included sequences from *B. fragilis* and *X. campestris*, suggesting this domain structure may appear widely across the Gram-negative bacteria (44). TonB7 and TonB11 both contain predicted CarboxypepD_reg-like domains (PF13715) which are often also found in TBDTs including SusC and the levan-targeting BT1763 as the N-terminal extension (NTE) (Fig 1) (27). The structure of this domain from BT1763 showed an Ig-like fold and deletion of this domain completely eliminated growth on levan (25). The association with both TBDT and TonB protein as well as the domain’s importance for BT1763 function may suggest a role in formation of the TBDT-TonB-ExbBD complex (25, 50). The DUF4488 domain (PF14869) found in TonB8 was structurally characterized in three *Bacteroides* proteins (51). The function of this domain is unclear but it appears to be restricted to the Bacteroidetes and the structures revealed an unknown ligand bound to each protein that suggests a role in binding small polar molecules such as carbohydrates (51). Most DUF4488 containing proteins are made up only of the DUF4488 domain but 19 of the 146 analyzed sequences contained a TonB C-terminal domain similar to TonB8 (51). TonB9 and 10 contain a second TonB C-terminal domain. This dual TonB domain structure seems to be limited to the Bacteroidetes (44). In both TonB9 and 10, the two TonB C-terminal domains share only a moderate sequence identity (62.8% and 73.1% respectively) that is similar or lower than the shared sequence identity of the domains of other *B. theta* TonB proteins (**Fig S1C,** shaded in blue).

Despite these differences in the full-length TonB proteins, the identified *B. theta* TonB C-terminal domains show moderate sequence similarity to the *E. coli* TonB C-terminal domain (37.8-49.4%) and high conservation of amino acids that are important for proper function of *E. coli* and *Pseudomonas aeruginosa* TonB (Fig 2, **Fig S1C**, **S2**). Complete conservation is seen at the YP motif (residues 163-164 in *E. coli* TonB) and at various points in the downstream region that forms the core of the domain. Notably, none of the conserved residues are in the final β-strand proposed to pair with TBDT TonBox (Fig 2B, C). The YP motif is the most conserved feature among TonB proteins (44). The tyrosine residue has been shown to interact with *E. coli* TBDTs BtuB and FecA and mutation of this residue results in a non-functional *P. aeruginosa* TonB1 (37, 52–54). The proline is not conserved in the second TonB domain of TonB9 but mutation of this residue in other TonB proteins does not disrupt function (54). Complete conservation is seen at residues equivalent to the *E. coli* TonB Gly 174, Gly186, and Trp213. The precise role of these residues is unclear but mutations in the Gly residues in *E. coli* TonB reduced *E. coli* growth on iron and sensitivity to colicins suggesting these residues are important for proper function of TonB (55). The equivalent residue to Gly174 in *P. aeruginosa* TonB1 (Gly275) is also essential for proper function (54). Although multimer formation by TonB is still unclear, Trp213 in *E. coli* TonB has been suggested to promote dimer formation as W213C mutants readily form cross-linked dimers (56). Conservation is also seen at *E. coli* Phe180 in all the sequences except Domain 2 of TonB10; however, this residue was not found to be highly conserved in a broader comparison of TonB proteins and it is unclear what role it plays in *E. coli* TonB (44). Finally, Val225 of *E. coli* TonB immediately precedes the region where β-sheet pairing occurs with the TonBox of TBDT but as side chains do not seem to be important for this interaction, it is unclear why this residue is conserved in all sequences analyzed except TonB11 where it is replaced with an isoleucine (Fig 2AB). Additionally, the corresponding valine in *P. aeruginosa* TonB1 (Val326) is well outside the β-strand that pairs with the TonBox of FoxA suggesting this residue may play an additional role in TonB function (Fig 2C).

To further understand the potential role of these eleven TonB proteins, we analyzed the genomic context of each of the genes (Fig 3). The *tonB* genes are dispersed throughout the genome, with most *tonB* genes being alone without other Ton complex genes. Notable exceptions to this are *tonB9* and *tonB10* which are found near each other, separated only by one gene predicted to encode a thioredoxin similar to DsbE. Additionally, *tonB5* (*bt2665*) is organized next to and in the same transcriptional orientation as predicted ExbB BT2668 and predicted ExbDs BT2666-2667. The *tonb4* (*bt2059*) gene is also found near predicted ExbB BT2055 and predicted ExbDs BT2052-2053 but the intervening genes include proteins of unknown function (hypothetical proteins), a hydrolase, and isoprenyl synthase that are not predicted to be involved in formation of the transport complex. Several *tonB* genes are found near transposases including *tonB5, tonB6, tonB9,* and *tonB10* although it is not clear if the *tonB* genes would be transferred by these transposases. Particularly interesting is *tonB8* which appears to be found at the end of the rhamnogalacturonan-II (RG-II) PUL 3 (20). The *bt3673* gene was previously annotated as a hypothetical protein and was not characterized as part of the previous exploration of RG-II degradation so it is not clear if this gene is important in RG-II degradation. Similarly*, tonB1, tonB2, tonB7, and tonB11* are found near predicted transcriptional regulators that may allow for better understanding of the control of expression of these genes.

**Figure 3.**
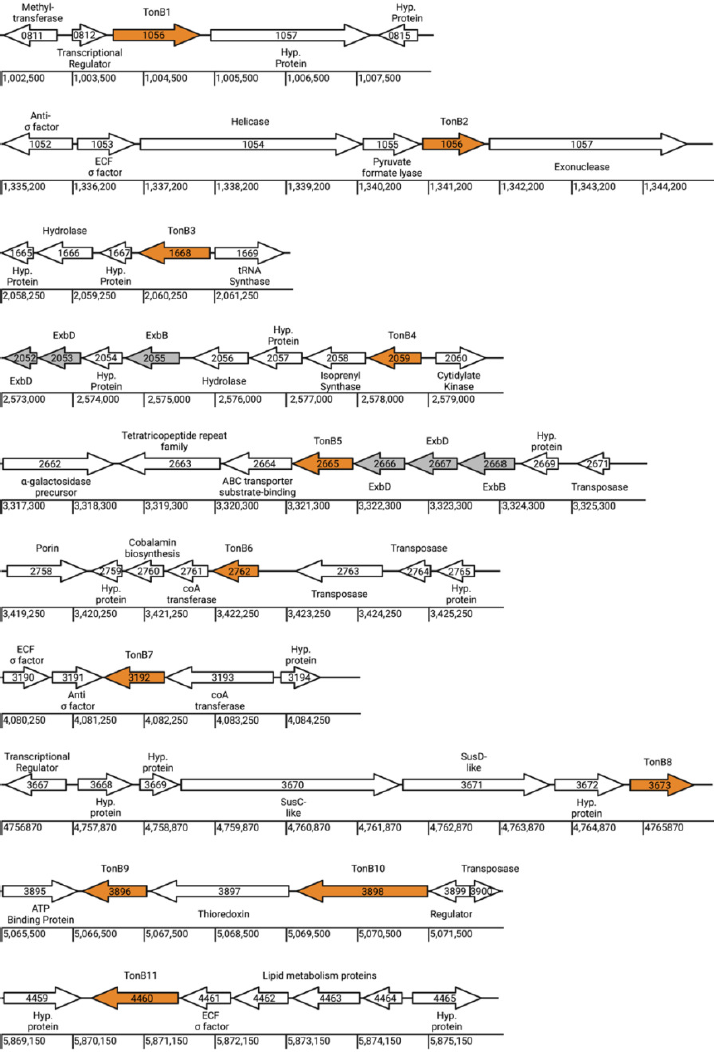
The genomic context of each of the eleven *tonB* genes. Each locus tag is shown within the arrow depicting the putative ORF. Arrow direction depicts the transcription orientation. 5cale is shown in base pairs using the reference sequence GCF_000011065.1. The predicted function of the peptide product is depicted outside each gene arrow. Hyp. protein: conserved hypothetical protein. Figure generated in BioRender.

Taken together, the conservation of key residues and the overall predicted C-terminal domains of the identified *B. theta* proteins suggest these proteins are capable of functioning as TonB proteins. However, the addition of unique domains to the overall protein architecture of several of these proteins may allow for formation of TBDT-TonB-ExbBD complexes that are functionally distinct from characterized complexes and the lack of genetic organization with ExbBD genes allows for unique assemblies of these complexes. To begin to explore the function of these TonB proteins, we first focused on the formation of a SusC-TonB pair by deleting both the TonBox of SusC and constructing in-frame deletions for each of the eleven *tonB* genes to explore the effect of each deletion on starch utilization.

### Deletion of the TonBox of SusC eliminates growth of *B. theta* on starch

The canonical *E. coli* TonBox consensus sequence is acidic-T-hydrophobic-hydrophobic-V-polar-A (26). Conservation of the canonical TonBox sequence is seen across many TBDTs but some divergence has made it difficult to confidently predict this motif from sequence alone. To identify the TonBox in SusC, we looked for two features: 1. High conservation across a short region preceding the putative plug of SusC-like transporters in *B. theta* and 2. Close alignments with the TonBox from characterized TBDTs from other bacteria.

Using an alignment of 100 SusC-like proteins from *B. theta* we identified a highly conserved region with the consensus sequence DEVVV(V/T/I) (representative sequences shown in Fig 4A, **S3**). This region also aligns well with the characterized TonBox sequences from FecA (52) and FhuE (57) from *E. coli*, HasR from *Serratia marcescens* (58), FoxA (34)and FpvA (59) from *P. aeruginosa,* and RagA from *P. gingivalis* (24) (Fig 4A, **S3**). Analysis of BT1763 from *B. theta* also identified this sequence as the TonBox and showed a significant change in function when the TonBox was mutated or deleted (25). Based on this, we propose that the TonBox sequence in SusC is DEVVVI found at residues 105 to 110.

**Figure 4.**
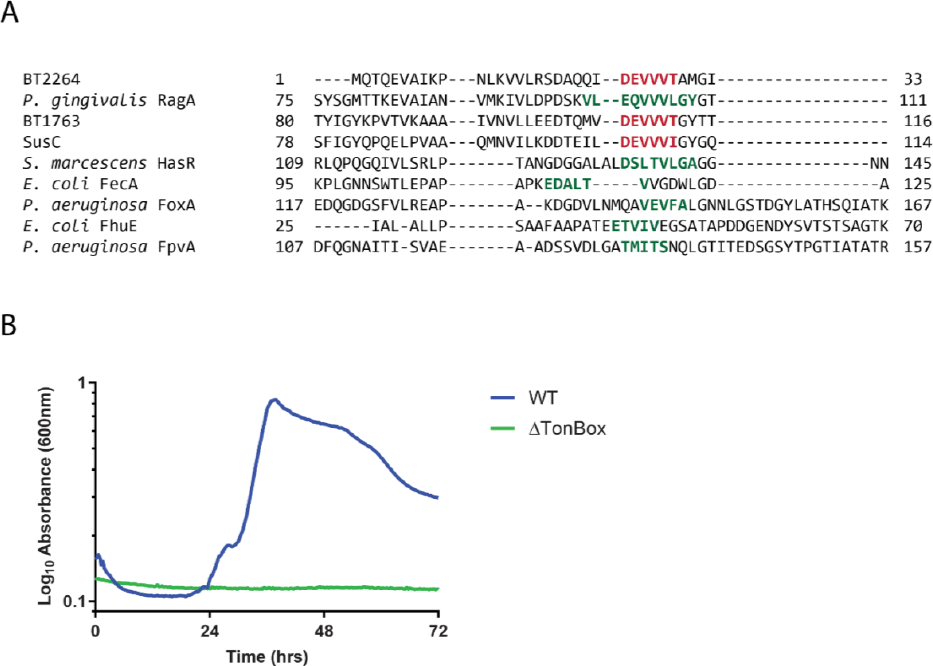
**A.** Excerpt of the multiple sequence alignment of 5usC from *B. thetaiotaomicron* with BT1763 and BT2264 for which structures have been determined and characterized TonB-dependent transporters from *Porphyromonas gingivalis, Escherichia coli*, *Serratia marcescens*, and *Pseudomonas aeruginosa*. Identified TonBoxes are shown in green, our proposed TonBoxes for the *B. thetaiotaomicron* transporters are shown in red. The full sequence alignment is shown in Fig. 53. **B.** Average growth curves of wild-type (WT) and the 5usC TonBox deletion (ΔTonBox) *B. thetaiotaomicron* cultured on 2.5 mg ml^−1^ potato amylopectin. Matched growth experiments in maltose and maltoheptataose are shown in Fig. 54.

We constructed an in-frame deletion of these six residues to create our ΔTonBox strain of *B. theta*. We chose to delete these residues rather than mutating them as previous studies have shown that mutations to chemically distinct residues often do not disrupt TBDT function but deletion of the TonBox disrupts function of the TBDT likely by eliminating pairing to TonB (26, 58). Our *B. theta*

ΔTonBox strain grows normally on maltose which does not have to be taken up through SusC (**Fig S4A**). However, the ΔTonBox strain cannot grow on amylopectin (Fig 4B) or other starch substrates including maltoheptaose (**Fig S4B**) that wild-type *B. theta* can efficiently utilize. This suggests that with the TonBox removed, SusC cannot pair with TonB to import these starch substrates supporting the role of SusC as a TonB-dependent transporter and the importance of this pairing. Interestingly, a similar TonBox deletion in the *B. theta* levan TBDT BT1763 caused only a lag in growth while a full deletion of the N-terminal extension (NTE) was needed to eliminate growth on levan (25). These results support the importance of the TonBox but suggest that further characterization of both the TonBox and NTE may be required to fully understand TBDT-TonB pairing across PUL.

### Deletion of TonB4 increases lag phase of *B. theta* growth on starch

We explored the role of each TonB by assessing the effect of these gene deletions on the function of the prototypical *Bacteroides* TBDT, SusC, during starch utilization. We began by using glucose or maltose as the sole carbon source as *B. theta* does not require TBDTs to import these sugars and therefore deletion of TonB proteins should not affect growth. However, deletion of TonB7, but no other TonB genes, resulted in consistently slower growth to both OD=0.3 and max OD on glucose (representative growth curves shown in Fig. 5A, growth time to OD=0.3 over four experiments shown in **Fig. S5A**) and maltose (**Fig S5B**). *B. theta* does require TBDTs to uptake vitamin B12 and heme which are found in the minimal media used for these growths so it is possible TonB7 pairs with at least one of these TBDTs; however, the TonB7 protein was not identified in previous proteomic analysis of *B. theta* conducted in similar media and well as the proteomic analysis presented here (60, 61). This suggests that the location of this gene in the *B. theta* genome may play a more important role than expression of TonB7 protein, requiring characterization beyond the scope of this work.

**Figure 5.**
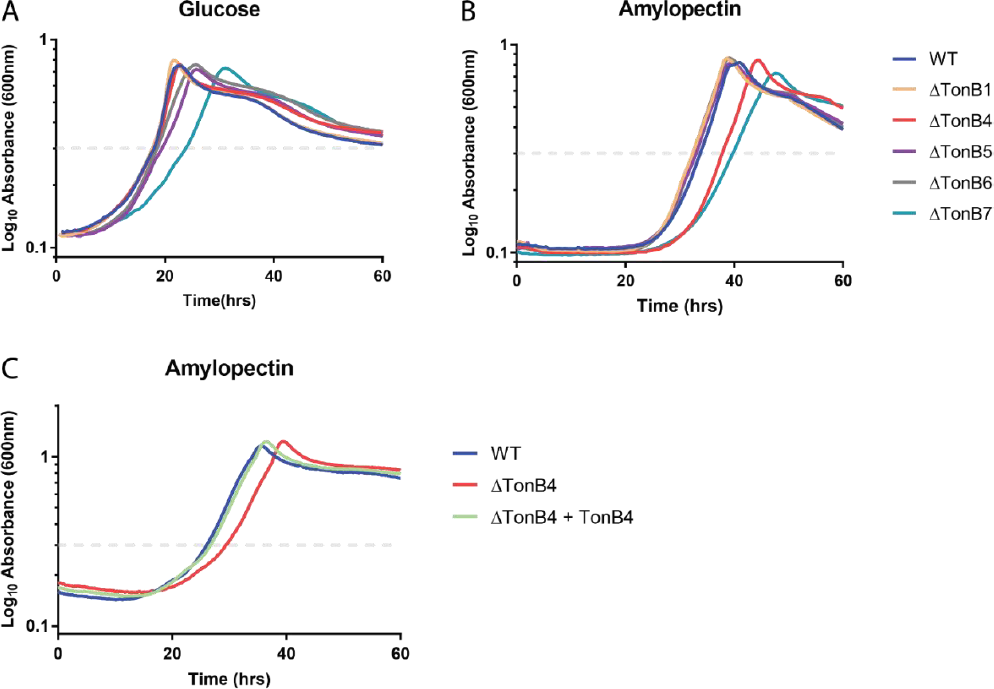
**A-B.** Representative average growth curves of wild-type (WT) and select TonB deletion strains of *B. thetaiotaomicron* cultured on 5 mg ml^−1^ glucose (A) and 2.5 mg ml^−1^ potato amylopectin (B}. The dash line indicates OD=0.3 which is used as a reference point for calculating lag time. Growth to OD=0.3 for all TonB deletions over four experiments and matched growth experiments in maltose are shown in Fig. 55A-D. **C.** Representative average growth curves of wild-type (WT}, TonB4 deletion strain, and the TonB4 deletion strain with the TonB4 gene complemented in another location on the genome cultured on 5 mg ml^−1^ potato amylopectin. Matched growth experiments in maltoheptaose and maize amylopectin are shown in Fig. 55E-F.

We next assessed growth on starch substrates. Deletion of TonB4 resulted in consistently slower growth to both OD=0.3 and max OD on potato amylopectin (representative growth curves shown in Fig 5B, growth time to OD=0.3 over four experiments shown in **S5C, D**). The slower growth of TonB7 was consistent with what was seen on glucose and maltose. The slower growth of TonB4 could be rescued by complementing the gene into another location on the chromosome suggesting that this reduced growth is due to the lack of TonB4 protein (Fig 5C, **Fig S5C, D**). Similar slow growth of the ΔTonB4 and a return to wild-type like growth in the complementation strain is also seen for other starch substrates including maltoheptaose (**Fig S5E**) and maize amylopectin (**Fig S5F**).

### TonB6 may compensate for loss of TonB4

That we observed a lag but not loss of growth for ΔTonB4 led us to question if there is a specific TonB protein that can replace TonB4 in pairing with SusC or if any of the remaining 10 TonB proteins could properly function with SusC. We have previously reported membrane proteomics to quantify amounts of Sus proteins in cells grown in the presence of maltose to induce Sus expression and TonB4 was the only TonB protein detected in those samples (60). We chose to revisit membrane proteomics for comparison with the ΔTonB4 strain using a tandem mass tag-based approach for peptide quantification between conditions and strains (**Table S1**). As expected, we saw a dramatic increase in Sus proteins when both WT and ΔTonB4 were grown on maltose as compared to glucose (SusC shown as an example in Fig 6 but similar results were seen for SusA-G).

**Figure 6.**
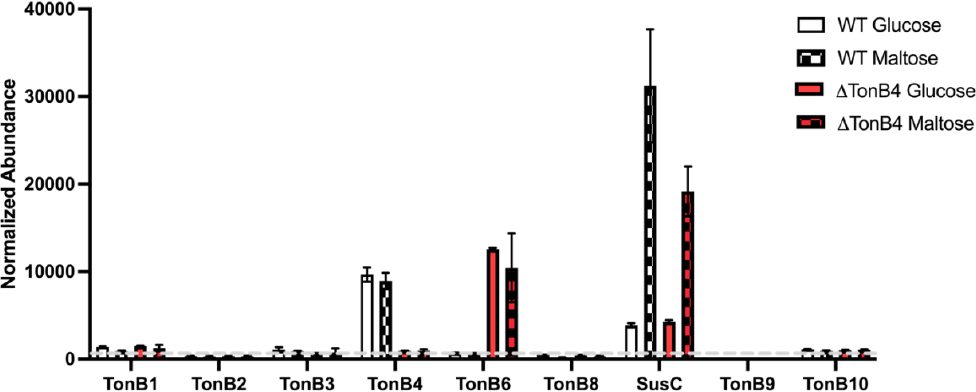
Normalized abundances of TonB and 5usC proteins from quantitative membrane proteomics. Mean and standard deviation of two samples are shown. Open bars show data from cells grown on 5 mg ml^−1^ glucose while hatched bars are from cells grown on 5 mg ml^−1^ maltose to induce 5us expression. White bars are samples isolated from wild-type (WT) cells while red bars are from the TonB4 deletion strain. The dashed line indicates background readings based on TonB4-matched peptides in the TonB4 deletion strain. Full data in Table 51. Quantitation was performed using high-quality M53 spectra, see Methods.

Like the previously published data, TonB4 was highly abundant in both the glucose and maltose grown WT cell membranes (Fig 6). Unlike the previous data, we also measured low amounts of other TonB proteins but notably did not see expression of TonB7 which also caused a growth defect when deleted. In the ΔTonB4 strain, abundance of most TonB proteins was unchanged; however, there was an apparent increase in TonB6. Interestingly, TonB6 appeared to be similarly abundant in the ΔTonB4 strain as TonB4 in the WT strain. Therefore, we hypothesize that TonB6 partially complements TonB4 in the ΔTonB4 strain. Furthermore, we have not been able to construct a ΔTonB4/6 double-deletion strain suggesting that the double mutant is lethal and that these TonB proteins play redundant roles.

### TonB4 shows a variable role in growth on other polysaccharides

This evidence supports a model where SusC is normally energized by the TonB4 protein, though it is unclear if this is a specific SusC-TonB4 interaction or if TonB4 is the preferred TonB for all polysaccharide utilization under normal lab conditions. To address this, we assessed the growth of the single TonB deletions on various polysaccharide substrates for which the PUL has been defined and it is known that a single TBDT is responsible for uptake including arabinan, levan, chondroitin sulfate, and pectic galactan (8, 14, 15, 25, 62). Interestingly, we see a variety of phenotypes across these four polysaccharides suggesting that the SusC-TonB4 interaction may not be unique, but TonB4 is also not the dominant TonB for all polysaccharide utilization (Fig 7). The ΔTonB4 strain shows slower growth to both OD=0.3 and max OD when grown on both pectic galactan and chondroitin sulfate (Fig 7A-B). This suggests that both TBDTs BT4671 (pectic galactan) and BT3332 (chondroitin sulfate) may primarily pair with TonB4 similarly to SusC. Additional work is needed to confirm if TonB6 is the secondary TonB for these transporters.

**Figure 7.**
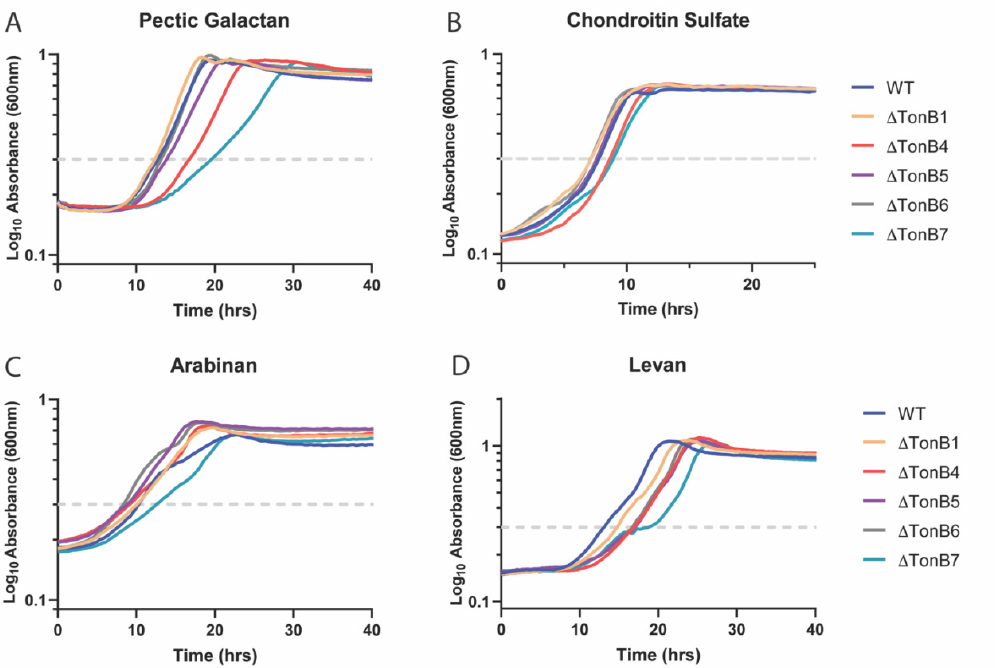
**A-D.** Representative average growth curves of wild-type (WT) and select TonB deletion strains of *B. thetaiotaomicron* cultured on 5 mg ml^−1^ pectic galactan (A}, 5 mg ml^−1^ chondroitin sulfate (B}, 5 mg ml^−1^ arabinan (C}, and 5 mg ml^−1^ levan (D}. The dash line indicates OD=0.3 which is used as a reference point for delayed growth.

Alternatively, the ΔTonB4 strain shows growth similar to other *B. theta* strains when grown on arabinan and levan suggesting that these transporters do not show a preference for pairing with TonB4 and multiple TonB proteins may be able to facilitate transport of these substrates with similar efficiency (Fig 7C-D).

### TonB genes vary across the Bacteroides genus

To understand conservation of the putative *B. theta* TonB proteins throughout the genus, we conducted a comparative genomics analysis by searching for homologs of each full-length *B. theta* TonB protein in a range of fully sequenced *Bacteroides* species (Fig 8, **Table S2 & S3**). We found that the set of TonB proteins in each species and even varying strains of the same species is highly divergent. Homologs of TonB4, TonB5, and TonB10 were found in almost all of the *Bacteroides* species we analyzed suggesting that these TonB proteins may play an essential role in *Bacteroides* physiology. This also supports that TonB4 may be widely important for polysaccharide uptake as seen in a recent work analyzing the TonB homologs in *B. fragilis* where the TonB4 homolog (*B. fragilis* TonB3) is essential for growth on a variety of polysaccharides (40). However, sequence similarity of these conserved proteins decreased in species less closely related to *B. theta*. This is particularly striking for TonB10 where many of the homologs do not consist of the same domain structure as the *B. theta* TonB10, resulting in a low overall sequence similarly. Some TonB proteins including homologs of TonB7 and TonB9 were found in only a few strains of *B. theta* (Fig 8, **Table S3**). Additional research is needed to understand the unique role these proteins are playing in these strains. Homologs of many TonB proteins such as TonB6, TonB8, and TonB11 are found only in species closely related to *B. theta*. This is particularly interesting in the case of TonB6 that is important for supplementing the function of TonB4 in *B. theta*. The lack of a TonB6 homolog suggests that many of these bacteria may show a higher dependence on proper function of the TonB4 homolog. Indeed, this was recently shown for *B. fragilis* 638R where deletion of the TonB4 homolog (*B. fragilis* TonB3) completely eliminates growth on polysaccharides and we were not able to identify a TonB6 homolog in this strain (40).

**Figure 8.**
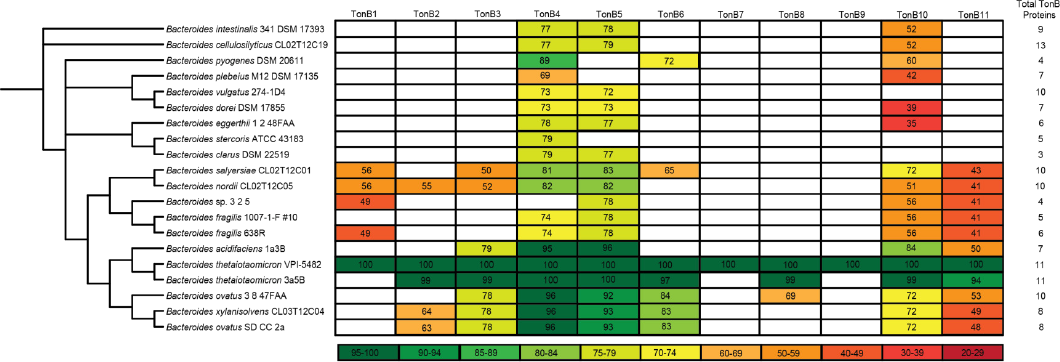
Percent sequence similarities between full-length *B. theta* TonB proteins (TonB1-11) and proteins in various Bacteroides genomes are shown in the table below. The phylogenetic tree shown on the left is based on 16s rRNA sequence similarity. The total TonB proteins identified in each genome is indicated to the right as many genomes contain predicted TonB proteins that do not show significant sequence similarity to the *B. theta* TonB proteins.

Many species of *Bacteroides* have additional TonB proteins that show little homology to the *B. theta* TonB proteins (**Table S4**). For example, *Bacteroides plebeius* contains TonB proteins with some homology to TonB4 and TonB10 but also contains 5 additional predicted TonB proteins (Fig 8, **Table S4**). Even in species closely related to *B. theta* such as *Bacteroides ovatus* and *Bacteroides acidifaciens*, we found predicted TonB proteins with little to no homology to the *B. theta* TonB proteins. While it is still unclear why the Bacteroides maintain such a large number of TonB proteins, the diversity of TonB proteins strongly suggest that they are important and further characterization of these proteins will allow us to better understand *Bacteroides* physiology.

## Discussion

By deleting either the TonBox portion of the *susC* gene or the *tonB4* gene, we provide data that supports that starch is taken up by *B. theta* through a TonB-dependent mechanism. While deletion of the *tonB4* gene causes slower bacterial growth, we show that levels of TonB6 proteins drastically increase in the absence of TonB4 suggesting that the TonB6 homolog can also energize transport of starch through SusC. The phenomenon does not seem to be restricted only to starch as growth on pectic galactan and chondroitin sulfate are similarly affected by the *tonB4* gene deletion. Interestingly growth on arabinan or levan is not affect by TonB deletion of any of the eleven TonB homologs. Taken together these results suggest that there is specificity of pairing between TBDT and TonB proteins in *B. theta* but that there is redundancy or overlapping function among some *B. theta* TonB homologs.

*B. theta* is often used as a model system to understand the *Bacteroides* but there are many unique aspects to each *Bacteroides* species and the TonB content of each species and even strain is no exception. Our comparative genomic analysis showed that while TonB4 is highly conserved across the *Bacteroides*, individual species typically contain an array of additional *tonB* genes that are often not highly conserved and may even be specialized for a limited number of strains within a single species.

This importance of TonB4 as well as the potential redundancy offered by TonB6 provide a useful explanation of previous data exploring the importance of various *B. theta* genes. In two separate transposon screens of *B. theta*, no TonB homologs were identified as essential genes (63, 64). However, the strain with a transposon insertion in the *tonB4* gene showed a decreased abundance after extended exponential growth and decreased abundance after mono-association in mice as compared to wild-type *B. theta (63)*. This suggests that the growth defect we see as lower growth in the ΔTonB4 strain persists in extended exponential growth and is sufficient to decrease the ability of this strain to colonize mice. However, likely due to the redundancy offered by TonB6, disruption of the *tonB4* gene does not eliminate growth or the ability to colonize the mouse intestine (63).

These differential growth outcomes must have a molecular basis in TBDT-TonB pair formation While we analyzed the role of the SusC TonBox in TBDT-TonB protein pairing, the TonBox is likely not the only region of interaction between these proteins (65). Because the predicted TonBox region is well conserved across Bacteroides TBDT and we see different specificity for TonB4 across the starch, arabinan, and levan transporters, it is likely that these other interactions are important for determining the specificity of pairing between TBDT and TonB proteins. Sequence variation in the N-terminus of the TBDT is likely important for this specificity although it is not currently clear if this is limited to the plug domain or if the N-terminal extension and signal transduction domains that are common in *Bacteroides* TBDT also play a role (27). It also seems likely that the sequence of the TonB protein is also highly important. Indeed, TonB4 and TonB6 show high sequence similarity and differences between these proteins may point to important regions for pairing specificity.

The transporter is also not the only protein that TonB must be in contact with to facilitate transport. TonB is also associated with the inner membrane proteins ExbB and ExbD. *B theta* contains 5 predicted ExbB homologs and 8 predicted ExbD homologs. Previous work in *E. coli* suggests that TonB interaction with ExbD is essential for TonB to adopt the correct confirmation for interaction with the TBDT (65, 66). Thus, it is likely that only some ExbB and ExbD homologs are capable of properly energizing TonB4 and TonB6 for the polysaccharide utilization explored in this paper. Exploration of this inner membrane complex is an essential component that must be explored to fully understand the requirements of TonB-dependent transport in the *Bacteroides*.

A significant open question is the role of the other nine TonB homologs in *B. theta*. While we have focused on polysaccharide transport here, it seems possible that other TonB homologs may be important for uptake of B12 and heme that are generally taken up by much smaller TBDTs although we did not see abundance of additional TonBs in the membrane proteomics (27). Additionally, it has been shown that for some substrates, TonB-like proteins may play an additional role in transport by directly interacting with the substrate (67). This provides a potential explanation for the unique domain structure of some of the TonB proteins characterized here. This is an important consideration as more TBDT substrates are identified and more TBDTs are characterized in *B. theta*.

While much focus has been given to carbohydrate-active enzymes and other outer-membrane proteins essential for polysaccharide utilization in the *Bacteroides*, this study extends our understanding of the larger protein complex required for polysaccharide utilization in *B. theta*. TonB-targeting drugs are currently being considered for pathogenic bacteria and a deeper understanding of this system in *B. theta* may offer new opportunities for manipulating both the microbiome and pathogenic *Bacteroides*. The importance and conservation of TonB4 suggests that drugs targeting this protein may offer a way to decrease growth of all *Bacteroides* while the redundancy offered by TonB6 may allow *B. theta* and related species to survive at low levels while species such a pathogenic *B. fragilis* are eliminated (40).

This work also highlights the many aspects left to understand about TonB-dependent transport. Along with the growing variety of substrates known to be transported through TBDT, the unique domain architectures seen in TonB proteins suggests the previously characterized structure-function relationships of the TBDT-TonB pair will not be sufficient to fully understand these systems.

## Materials and Methods

### Bacterial strains and culture conditions

The *B. thetaiotaomicron* VPI-5482 Δtdk strain is the parent strain for all mutations used in this study and is referred to as wild type (WT). Mutant strains were generated via allelic exchange as previously described (18, 68). Briefly, the genomic region containing the desired gene deletion was inserted into the counter selectable allelic exchange vector pExchange_tdk. The primers used in this study were synthesized by IDT DNA Technologies and are described in **Table S5**. A summary of all plasmids and strains used in this study is provided in **Table S6**.

All *B. theta* strains were cultured in a 37°C Coy anaerobic chamber (5% H_2_/10% CO_2_/85% N_2_) from freezer stocks into tryptone-yeast extract-glucose (TYG) medium (69) and grown for 16 h to an O.D._600_ ∼ 1.0. The cells were then back diluted 1:100 into Bacteroides minimal media (MM) including 5 mg/ml glucose and grown overnight (16 h).

For kinetic growth experiments in a plate reader, the MM-glucose grown cells were then washed in minimal media containing no carbon and back diluted approximately 1:200 into MM with the experimental carbohydrate, glucose, or maltose to a final volume of 200uL. Thus, both glucose and maltose controls and the experimental carbohydrate grown cultures were started at the same initial O.D._600_ of 0.1. The carbon sources used for comparison to glucose-grown cultures included: 5 mg/ml maltose (Sigma), 2.5 mg/ml (2.17 mM) maltoheptaose (Carbosynth), and 5 mg/ml potato amylopectin (Sigma). Growths were conducted in a flat bottom 96-well plate (Costar) covered with a gas permeable, optically clear polyurethane membrane (Diversified Biotech, USA). Plates were loaded in a Biostack automated plate-handling device (Biotek Instruments, USA) coupled with a Powerwave HT absorbance reader (Biotek Instruments, USA) inside the anaerobic chamber and O.D._600_ was recorded every 10-30 min. All plate reader growth experiments were performed in triplicate unless otherwise noted and the averages are reported in each figure. All biological experiments were repeated at least twice to verify consistent growth phenotypes from day to day.

### Gene complementation

The *tonB4* (*bt2059*) gene in a pNBU2 vector containing a constitutively active promoter was introduce into the genome of the ΔTonB4 *B. theta* strain in a single copy as previously described (16, 70). Briefly, the *bt2059* gene was introduced into the pNBU2_erm_us1311 plasmid using restriction enzyme cloning and primers in **Table S5**. After conjugative transfer of the plasmid into the ΔTonB4 *B. theta* strain, the plasmid integrated into the genome at the NBU2 *att2* site.

### Membrane Proteomics

#### Sample preparation

All strains were cultured in TYG and back diluted into MM containing glucose as described above. The MM-glucose grown cells were then back diluted 1:100 into 50mL of MM containing 5mg/ml glucose or maltose as indicated. The O.D._600_ was monitored every 30-45 minutes and the cells were harvested at mid-log (O.D. ∼0.7-0.8) by centrifugation at 5000 xg for 5 min. The cell pellet was frozen in liquid nitrogen and stored at −80°C.

To prepare the membrane faction, cells were thawed and resuspended in 1mL of 20mM KH_2_PO_4_ pH 7.3. The slurry was gently sonicated on ice. Intact cells were removed by centrifugation at 13,000 xg for 10 minutes at 4°C. The remaining soluble fraction was ultracentrifuged at 200,000 xg for 2 hrs at 4°C to pellet total membranes. The supernatant was removed and the membrane pellet resuspended in the same buffer, followed by a second round of ultracentrifugation at 200,000 xg. The resulting membrane pellet was resuspended in 20mM KH_2_PO_4_, 0.1% Tween-20 pH 7.3. Total protein in the final sample was quantified using the BCA Protein Assay Kit (Pierce).

The total membrane samples were submitted to the Mass Spectrometry-Based Proteomics Resource Facility in the Department of Pathology at the University of Michigan (Ann Arbor, MI). Samples were then processed using the TMT 10-plex Mass Tag Labeling Kit (Thermo Scientific) similar to manufacturer’s protocol and as previously adapted (71). Briefly, upon reduction (5 mM DTT, for 30 min at 45°C) and alkylation of cysteines (15 mM 2-chloroacetamide, for 30 min at room temperature), the proteins were precipitated by adding 6 volumes of ice-cold acetone followed by overnight incubation at −20°C. The precipitate was spun down, and the pellet was allowed to air dry. The pellet was resuspended in 0.1M TEAB and overnight (∼16 h) digestion with trypsin/Lys-C mix (1:25 protease:protein; Promega) at 37° C was performed with constant mixing using a thermomixer. The TMT 10-plex reagents were dissolved in 41 μl of anhydrous acetonitrile and labeling was performed by transferring the entire digest to TMT reagent vial and incubating at room temperature for 1 h. Reaction was quenched by adding 8 μl of 5% hydroxyl amine and further 15 min incubation. Labeled samples were mixed together, and dried using a vacufuge. An offline fractionation of the combined sample (∼200 μg) into 8 fractions was performed using high pH reversed-phase peptide fractionation kit according to the manufacturer’s protocol (Pierce; Cat #84868). Fractions were dried and reconstituted in 9 μl of 0.1% formic acid/2% acetonitrile in preparation for LC-MS/MS analysis. Details on sample preparation as well as the sample-to-TMT channel are found in Table S1.

#### Liquid chromatography-mass spectrometry analysis (LC-multinotch MS3)

In order to obtain superior quantitation accuracy, we employed multinotch-MS3 which minimizes the reporter ion ratio distortion resulting from fragmentation of co-isolated peptides during MS analysis (72). Orbitrap Fusion (Thermo Fisher Scientific) and RSLC Ultimate 3000 nano-UPLC (Dionex) was used to acquire the data. Two μl of the sample was resolved on a PepMap RSLC C18 column (75 μm i.d. x 50 cm; Thermo Scientific) at the flow-rate of 300 nl/min using 0.1% formic acid/acetonitrile gradient system (2-22% acetonitrile in 150 min;22-32% acetonitrile in 40 min; 20 min wash at 90% followed by 50 min re-equilibration) and directly sprayed onto the mass spectrometer using EasySpray source (Thermo Fisher Scientific). Mass spectrometer was set to collect one MS1 scan (Orbitrap; 120K resolution; AGC target 2×10^5^; max IT 100 ms) followed by data-dependent, “Top Speed” (3 seconds) MS2 scans (collision induced dissociation; ion trap; NCE 35; AGC 5×10^3^; max IT 100 ms). For multinotch-MS3, top 10 precursors from each MS2 were fragmented by HCD followed by Orbitrap analysis (NCE 55; 60K resolution; AGC 5×10^4^; max IT 120 ms, 100-500 m/z scan range).

#### Data analysis

Proteome Discoverer (v2.4; Thermo Fisher) was used for data analysis. MS2 spectra were searched against SwissProt *Bacteroides thetaiotaomicron* VPI-5482 (ATCC strain 29148) protein database using the following search parameters: MS1 and MS2 tolerance were set to 10 ppm and 0.6 Da, respectively; carbamidomethylation of cysteines (57.02146 Da) and TMT labeling of lysine and N-termini of peptides (229.16293 Da) were considered static modifications; oxidation of methionine (15.9949 Da) and deamidation of asparagine and glutamine (0.98401 Da) were considered variable. Identified proteins and peptides were filtered to retain only those that passed ≤1% FDR threshold. Quantitation was performed using high-quality MS3 spectra (Average signal-to-noise ratio of 10 and <50% isolation interference).

The mass spectrometry proteomics data have been deposited to the ProteomeXchange Consortium via the PRIDE (73) partner repository with the dataset identifier PXD041518.

### Protein Sequence Analysis

Protein domains including the TonB protein C-terminal domains were identified using Pfam version 32.0 (74). To identify the eleven potential TonB proteins, the complete genome of *Bacteroides thetaiotaomicron* VPI-5482 was searched for sequences that matched to PF03544 using the Joint Genome Institute’s Integrated Microbial Genomes & Microbiomes database (75). Each sequence was then searched for additional Pfam domains using the sequence search on the EMBL-EBI Pfam database (76). Predictions of transmembrane helices were made using the TMHMM Server v2.0 (77, 78) and signal peptides were predicted using SignalP-5.0 (79).

Multiple sequence alignment of TonB-dependent transporters and TonB proteins were conducted in Clustal Omega (76). Sequence similarity between TonB proteins was determined using the EMBOSS Needle pairwise sequence alignment (76).

### Genomic Context Analysis

The genomic context of each *tonB* gene was identified by browsing the complete genome of *Bacteroides thetaiotaomicron* VPI-5482 (GCF_000011065.1). Size, direction, and location of the surrounding genes were annotated, generally included all genes that are encoded on the same strand or all genes that appear to be co-transcribed. Protein function predictions were also analyzed using UniProt Release 2023_02 and the most informative prediction between the two was used (80).

### TonB Homology Analysis

To identify homologues of the 11 TonB proteins found in *B. theta*, we searched the Integrated Microbial Genomes (IMG) database (current as of May 2018) for all *Bacteroides* genome sequences and performed BLAST searches of each TonB protein with an E-value cutoff of 1e-50. We chose this stringent cutoff to limit the number of homologues that would match to several of our TonB proteins of interest. These results are shown in Table S2. Despite using this stringent cutoff, we still found that many TonB proteins in other *Bacteroides* genomes matched to several *B. theta* TonB proteins. For each genome in our dataset, we used the E-value and bit score generated through the BLAST search to match each TonB to the single *B. theta* TonB protein with the highest match. These full results are shown in Table S3 and select genomes are shown in Figure 8.

Additional TonB proteins in each *Bacteroides* genome were identified by searching for proteins with regions matching to the conserved protein domain family Pfam 03544: Gram-negative bacterial TonB protein C-terminal. The full list of matches to Pfam03544 are shown in Table S4 and totals for select genomes are shown in Figure 8. Phylogenetic tree in Figure 8 was constructed using the 16s rRNA gene from each strain shown.

### Protein Structure Visualization

Structures of E. coli TonB and Pseudomonas aeruginosa TonB1 were visualized in PyMOL (81).

## Acknowledgements

This work was supported by a Research Diversity Supplement to grant R01GM118475 awarded to R.M.P. and N.M.K.

We thank Dr. Venkatesha Basrur for assistance with the membrane proteomics work and all members of the Koropatkin and Martens labs for useful feedback and technical assistance.

## References

1. Salyers AA, West SE, Vercellotti JR, Wilkins TD. 1977. Fermentation of mucins and plant polysaccharides by anaerobic bacteria from the human colon. Applied and Environmental Microbiology 34:529–533.

2. J. Cummings. 1981. Short chain fatty acids in the human colon. Gut 22:763–79.

3. Wong JMW, de Souza R, Kendall CWC, Emam A, Jenkins DJA. 2006. Colonic Health: Fermentation and Short Chain Fatty Acids: Journal of Clinical Gastroenterology 40:235–243.

4. Hamer HM, Jonkers D, Venema K, Vanhoutvin S, Troost FJ, Brummer R-J. 2007. The role of butyrate on colonic function: Alimentary Pharmacology & Therapeutics 27:104–119.

5. Sonnenburg JL, Xu J, Leip DD, Chen C-H, Westover BP, Weatherford J, Buhler JD, Gordon JI. 2005. Glycan Foraging in Vivo by an Intestine-Adapted Bacterial Symbiont. Science 307:1955–1959.

6. El Kaoutari A, Armougom F, Gordon JI, Raoult D, Henrissat B. 2013. The abundance and variety of carbohydrate-active enzymes in the human gut microbiota. Nature Reviews Microbiology 11:497– 504.

7. Martens EC, Koropatkin NM, Smith TJ, Gordon JI. 2009. Complex Glycan Catabolism by the Human Gut Microbiota: The Bacteroidetes Sus-like Paradigm. J Biol Chem 284:24673–24677.

8. Martens EC, Lowe EC, Chiang H, Pudlo NA, Wu M, McNulty NP, Abbott DW, Henrissat B, Gilbert HJ, Bolam DN, Gordon JI. 2011. Recognition and Degradation of Plant Cell Wall Polysaccharides by Two Human Gut Symbionts. PLoS Biol 9:e1001221.

9. McNulty NP, Wu M, Erickson AR, Pan C, Erickson BK, Martens EC, Pudlo NA, Muegge BD, Henrissat B, Hettich RL, Gordon JI. 2013. Effects of Diet on Resource Utilization by a Model Human Gut Microbiota Containing Bacteroides cellulosilyticus WH2, a Symbiont with an Extensive Glycobiome. PLoS Biol 11:e1001637.

10. Grondin JM, Tamura K, Déjean G, Abbott DW, Brumer H. 2017. Polysaccharide Utilization Loci: Fueling Microbial Communities. J Bacteriol 199:e00860–16, e00860-16.

11. Anderson KL, Salyers AA. 1989. Biochemical evidence that starch breakdown by Bacteroides thetaiotaomicron involves outer membrane starch-binding sites and periplasmic starch-degrading enzymes. J Bacteriol 171:3192–3198.

12. Anderson KL, Salyers AA. 1989. Genetic evidence that outer membrane binding of starch is required for starch utilization by Bacteroides thetaiotaomicron. Journal of Bacteriology 171:3199– 3204.

13. Xu J, Bjursell MK, Himrod J, Deng S, Carmichael LK, Chiang HC, Hooper LV, Gordon JI. 2003. A genomic view of the human-Bacteroides thetaiotaomicron symbiosis. Science 299:2074–2076.

14. Luis AS, Briggs J, Zhang X, Farnell B, Ndeh D, Labourel A, Baslé A, Cartmell A, Terrapon N, Stott K, Lowe EC, McLean R, Shearer K, Schückel J, Venditto I, Ralet M-C, Henrissat B, Martens EC, Mosimann SC, Abbott DW, Gilbert HJ. 2018. Dietary pectic glycans are degraded by coordinated enzyme pathways in human colonic Bacteroides. Nat Microbiol 3:210–219.

15. Sonnenburg ED, Zheng H, Joglekar P, Higginbottom SK, Firbank SJ, Bolam DN, Sonnenburg JL. 2010. Specificity of Polysaccharide Use in Intestinal Bacteroides Species Determines Diet-Induced Microbiota Alterations. Cell 141:1241–1252.

16. Martens EC, Chiang HC, Gordon JI. 2008. Mucosal Glycan Foraging Enhances Fitness and Transmission of a Saccharolytic Human Gut Bacterial Symbiont. Cell Host & Microbe 4:447–457.

17. Cartmell A, Lowe EC, Baslé A, Firbank SJ, Ndeh DA, Murray H, Terrapon N, Lombard V, Henrissat B, Turnbull JE, Czjzek M, Gilbert HJ, Bolam DN. 2017. How members of the human gut microbiota overcome the sulfation problem posed by glycosaminoglycans. Proc Natl Acad Sci USA 114:7037– 7042.

18. Koropatkin NM, Martens EC, Gordon JI, Smith TJ. 2008. Starch catabolism by a prominent human gut symbiont is directed by the recognition of amylose helices. Structure 16:1105–1115.

19. Cuskin F, Lowe EC, Temple MJ, Zhu Y, Cameron E, Pudlo NA, Porter NT, Urs K, Thompson AJ, Cartmell A, Rogowski A, Hamilton BS, Chen R, Tolbert TJ, Piens K, Bracke D, Vervecken W, Hakki Z, Speciale G, Munōz-Munōz JL, Day A, Peña MJ, McLean R, Suits MD, Boraston AB, Atherly T, Ziemer CJ, Williams SJ, Davies GJ, Abbott DW, Martens EC, Gilbert HJ. 2015. Human gut Bacteroidetes can utilize yeast mannan through a selfish mechanism. Nature 517:165–169.

20. Ndeh D, Rogowski A, Cartmell A, Luis AS, Baslé A, Gray J, Venditto I, Briggs J, Zhang X, Labourel A, Terrapon N, Buffetto F, Nepogodiev S, Xiao Y, Field RA, Zhu Y, O’Neil MA, Urbanowicz BR, York WS, Davies GJ, Abbott DW, Ralet M-C, Martens EC, Henrissat B, Gilbert HJ. 2017. Complex pectin metabolism by gut bacteria reveals novel catalytic functions. Nature 544:65–70.

21. White JBR, Silale A, Feasey M, Heunis T, Zhu Y, Zheng H, Gajbhiye A, Firbank S, Baslé A, Trost M, Bolam DN, van den Berg B, Ranson NA. 2023. Outer membrane utilisomes mediate glycan uptake in gut Bacteroidetes. Nature 1–7.

22. Reeves AR, Wang GR, Salyers AA. 1997. Characterization of four outer membrane proteins that play a role in utilization of starch by Bacteroides thetaiotaomicron. J Bacteriol 179:643–649.

23. Glenwright AJ, Pothula KR, Bhamidimarri SP, Chorev DS, Baslé A, Firbank SJ, Zheng H, Robinson CV, Winterhalter M, Kleinekathöfer U, Bolam DN, van den Berg B. 2017. Structural basis for nutrient acquisition by dominant members of the human gut microbiota. Nature 541:407–411.

24. Madej M, White JBR, Nowakowska Z, Rawson S, Scavenius C, Enghild JJ, Bereta GP, Pothula K, Kleinekathoefer U, Baslé A, Ranson NA, Potempa J, van den Berg B. 2020. Structural and functional insights into oligopeptide acquisition by the RagAB transporter from Porphyromonas gingivalis. Nat Microbiol https://doi.org/10.1038/s41564-020-0716-y.

25. Gray DA, White JBR, Oluwole AO, Rath P, Glenwright AJ, Mazur A, Zahn M, Baslé A, Morland C, Evans SL, Cartmell A, Robinson CV, Hiller S, Ranson NA, Bolam DN, van den Berg B. 2021. Insights into SusCD-mediated glycan import by a prominent gut symbiont. 1. Nature Communications 12:1–14.

26. Kadner RJ. 1990. Vitamin B12 transport in Escherichia coli: energy coupling between membranes. Mol Microbiol 4:2027–2033.

27. Pollet RM, Martin LM, Koropatkin NM. 2021. TonB-dependent transporters in the Bacteroidetes: Unique domain structures and potential functions. Molecular Microbiology 115:490–501.

28. Celia H, Botos I, Ni X, Fox T, De Val N, Lloubes R, Jiang J, Buchanan SK. 2019. Cryo-EM structure of the bacterial Ton motor subcomplex ExbB–ExbD provides information on structure and stoichiometry. Commun Biol 2:358.

29. Biou V, Adaixo RJD, Chami M, Coureux P-D, Laurent B, Enguéné VYN, de Amorim GC, Izadi-Pruneyre N, Malosse C, Chamot-Rooke J, Stahlberg H, Delepelaire P. 2022. Structural and molecular determinants for the interaction of ExbB from Serratia marcescens and HasB, a TonB paralog. Commun Biol 5:355.

30. Celia H, Noinaj N, Zakharov SD, Bordignon E, Botos I, Santamaria M, Barnard TJ, Cramer WA, Lloubes R, Buchanan SK. 2016. Structural insight into the role of the Ton complex in energy transduction. Nature 538:60–65.

31. Sverzhinsky A, Chung JW, Deme JC, Fabre L, Levey KT, Plesa M, Carter DM, Lypaczewski P, Coulton JW. 2015. Membrane Protein Complex ExbB _4_-ExbD _1_-TonB _1_ from Escherichia coli Demonstrates Conformational Plasticity. J Bacteriol 197:1873–1885.

32. Higgs PI, Larsen RA, Postle K. 2002. Quantification of known components of the Escherichia coli TonB energy transduction system: TonB, ExbB, ExbD and FepA: TonB, ExbB, ExbD and FepA ratios. Molecular Microbiology 44:271–281.

33. Domingo Köhler S, Weber A, Howard SP, Welte W, Drescher M. 2010. The proline-rich domain of TonB possesses an extended polyproline II-like conformation of sufficient length to span the periplasm of Gram-negative bacteria. Protein Science 19:625–630.

34. Josts I, Veith K, Tidow H. 2019. Ternary structure of the outer membrane transporter FoxA with resolved signalling domain provides insights into TonB-mediated siderophore uptake. eLife 8:e48528.

35. Celia H, Noinaj N, Buchanan SK. 2020. Structure and Stoichiometry of the Ton Molecular Motor. IJMS 21:375.

36. Pawelek PD, Croteau N, Ng-Thow-Hing C, Khursigara CM, Moiseeva N, Allaire M, Coulton JW. 2006. Structure of TonB in Complex with FhuA, E. coli Outer Membrane Receptor. Science 312:1399– 1402.

37. Shultis DD, Purdy MD, Banchs CN, Wiener MC. 2006. Outer Membrane Active Transport: Structure of the BtuB:TonB Complex. Science 312:1396–1399.

38. Phansopa C, Roy S, Rafferty JB, Douglas CWI, Pandhal J, Wright PC, Kelly DJ, Stafford GP. 2014. Structural and functional characterization of NanU, a novel high-affinity sialic acid-inducible binding protein of oral and gut-dwelling Bacteroidetes species. Biochemical Journal 458:499–511.

39. Tauzin AS, Laville E, Xiao Y, Nouaille S, Le Bourgeois P, Heux S, Portais J-C, Monsan P, Martens EC, Potocki-Veronese G, Bordes F. 2016. Functional characterization of a gene locus from an uncultured gut *Bacteroides* conferring xylo-oligosaccharides utilization to *Escherichia coli*: Carbohydrate transporters of gut bacteria. Molecular Microbiology 102:579–592.

40. Parker AC, Seals NL, Baccanale CL, Rocha ER. 2022. Analysis of six tonB gene homologs in Bacteroides fragilis revealed that tonB3 is essential for survival in experimental intestinal colonization and intra-abdominal infection. Infection and Immunity 90:e00469–21.

41. Blanvillain S, Meyer D, Boulanger A, Lautier M, Guynet C, Denancé N, Vasse J, Lauber E, Arlat M. 2007. Plant Carbohydrate Scavenging through TonB-Dependent Receptors: A Feature Shared by Phytopathogenic and Aquatic Bacteria. PLoS ONE 2:e224.

42. Lohmiller S, Hantke K, Patzer SI, Braun V. 2008. TonB-dependent maltose transport by Caulobacter crescentus. Microbiology 154:1748–1754.

43. Liao H, Cheng X, Zhu D, Wang M, Jia R, Chen S, Chen X, Biville F, Liu M, Cheng A. 2015. TonB Energy Transduction Systems of Riemerella anatipestifer Are Required for Iron and Hemin Utilization. PLoS ONE 10:e0127506.

44. Chu BCH, Peacock RS, Vogel HJ. 2007. Bioinformatic analysis of the TonB protein family. Biometals 20:467–483.

45. Larsen RA, Wood GE, Postle K. 1993. The conserved proline-rich Motif is not essential for energy transduction by Escherichia coliTonB protein. Mol Microbiol 10:943–953.

46. Evans JS, Levine BA, Trayer IP, Dorman CJ, Higgins CF. 1986. Sequence-imposed structural constraints in the TonB protein of E. coli. FEBS Letters 208:211–216.

47. Virtanen SI, Kiirikki AM, Mikula KM, Iwaï H, Samuli Ollila OH. 2020. Heterogeneous dynamics in partially disordered proteins. Physical Chemistry Chemical Physics 22:21185–21196.

48. Zhang HZ, Hackbarth CJ, Chansky KM, Chambers HF. 2001. A Proteolytic Transmembrane Signaling Pathway and Resistance to β-Lactams in Staphylococci. Science 291:1962–1965.

49. Rasmussen BA, Bush K, Tally FP. 1993. Antimicrobial Resistance in Bacteroides. Clinical Infectious Diseases 16 (Suppl 4):S390–400.

50. Bolam DN, van den Berg B. 2018. TonB-dependent transport by the gut microbiota: novel aspects of an old problem. Current Opinion in Structural Biology 51:35–43.

51. Kumar A, Punta M, Axelrod HL, Das D, Farr CL, Grant JC, Chiu H-J, Miller MD, Coggill PC, Klock HE, Elsliger M-A, Deacon AM, Godzik A, Lesley SA, Wilson IA. 2014. Crystal structures of three representatives of a new Pfam family PF14869 (DUF4488) suggest they function in sugar binding/uptake: Crystal Structures of Pfam PF14869 (DUF4488). Protein Science 23:1380–1391.

52. Ogierman M, Braun V. 2003. Interactions between the Outer Membrane Ferric Citrate Transporter FecA and TonB: Studies of the FecA TonB Box. JB 185:1870–1885.

53. Cadieux N, Kadner RJ. 1999. Site-directed disulfide bonding reveals an interaction site between energy-coupling protein TonB and BtuB, the outer membrane cobalamin transporter. Proceedings of the National Academy of Sciences 96:10673–10678.

54. Zhao Q, Poole K. 2002. Mutational Analysis of the TonB1 Energy Coupler of Pseudomonas aeruginosa. JB 184:1503–1513.

55. Traub I, Gaisser S, Braun V. 1993. Activity domains of the TonB protein. Mol Microbiol 8:409–423.

56. Vakharia-Rao H, Kastead KA, Savenkova MI, Bulathsinghala CM, Postle K. 2007. Deletion and Substitution Analysis of the Escherichia coli TonB Q160 Region. JB 189:4662–4670.

57. Sauer M, Hantke K, Braun V. 1990. Sequence of the fhuE outer-membrane receptor gene of Escherichia coli K12 and properties of mutants. Mol Microbiol 4:427–437.

58. Lefèvre J, Delepelaire P, Delepierre M, Izadi-Pruneyre N. 2008. Modulation by Substrates of the Interaction between the HasR Outer Membrane Receptor and Its Specific TonB-like Protein, HasB. Journal of Molecular Biology 378:840–851.

59. Cobessi D, Celia H, Folschweiller N, Schalk IJ, Abdallah MA, Pattus F. 2005. The Crystal Structure of the Pyoverdine Outer Membrane Receptor FpvA from Pseudomonas aeruginosa at 3.6Å Resolution. Journal of Molecular Biology 347:121–134.

60. Tuson HH, Foley MH, Koropatkin NM, Biteen JS. 2018. The Starch Utilization System Assembles around Stationary Starch-Binding Proteins. Biophysical Journal 115:242–250.

61. Valguarnera E, Scott NE, Azimzadeh P, Feldman MF. 2018. Surface Exposure and Packing of Lipoproteins into Outer Membrane Vesicles Are Coupled Processes in *Bacteroides*. mSphere 3:e00559–18, /msphere/3/6/mSphere559-18.atom.

62. Raghavan V, Groisman EA. 2015. Species-Specific Dynamic Responses of Gut Bacteria to a Mammalian Glycan. J Bacteriol 197:1538–1548.

63. Goodman AL, McNulty NP, Zhao Y, Leip D, Mitra RD, Lozupone CA, Knight R, Gordon JI. 2009. Identifying Genetic Determinants Needed to Establish a Human Gut Symbiont in Its Habitat. Cell Host & Microbe 6:279–289.

64. Liu H, Shiver AL, Price MN, Carlson HK, Trotter VV, Chen Y, Escalante V, Ray J, Hern KE, Petzold CJ, Turnbaugh PJ, Huang KC, Arkin AP, Deutschbauer AM. 2021. Functional genetics of human gut commensal Bacteroides thetaiotaomicron reveals metabolic requirements for growth across environments. Cell Rep 34:108789.

65. Gresock MG, Postle K. 2017. Going Outside the TonB Box: Identification of Novel FepA-TonB Interactions In Vivo. J Bacteriol 199:e00649–16.

66. Gresock MG, Kastead KA, Postle K. 2015. From Homodimer to Heterodimer and Back: Elucidating the TonB Energy Transduction Cycle. J Bacteriol 197:3433–3445.

67. Wojnowska M, Walker D. 2020. FusB Energizes Import across the Outer Membrane through Direct Interaction with Its Ferredoxin Substrate. 5. mBio 11.

68. Cameron EA, Maynard MA, Smith CJ, Smith TJ, Koropatkin NM, Martens EC. 2012. Multidomain Carbohydrate-binding Proteins Involved in Bacteroides thetaiotaomicron Starch Metabolism. J Biol Chem 287:34614–34625.

69. Holdeman LV, Cato EP, Moore W. 1977. Anaerobe Laboratory Manual, 4th ed. V.P.I. Anaerobe Laboratory, Virginia Polytechnic Institute and State University, Blacksburg, Virginia.

70. Degnan PH, Barry NA, Mok KC, Taga ME, Goodman AL. 2014. Human gut microbes use multiple transporters to distinguish vitamin B_12_ analogs and compete in the gut. Cell Host Microbe 15:47–57.

71. Tank EM, Figueroa-Romero C, Hinder LM, Bedi K, Archbold HC, Li X, Weskamp K, Safren N, Paez-Colasante X, Pacut C, Thumma S, Paulsen MT, Guo K, Hur J, Ljungman M, Feldman EL, Barmada SJ. 2018. Abnormal RNA stability in amyotrophic lateral sclerosis. Nat Commun 9:2845.

72. McAlister GC, Nusinow DP, Jedrychowski MP, Wϋhr M, Huttlin E L, Erickson BK, Rad R, Haas W, Gygi SP. 2014. MultiNotch MS3 enables accurate, sensitive, and multiplexed detection of differential expression across cancer cell line proteomes. Anal Chem 86:7150–7158.

73. Perez-Riverol Y, Bai J, Bandla C, García-Seisdedos D, Hewapathirana S, Kamatchinathan S, Kundu Perez-Riverol Y, Bai J, Bandla C, García-Seisdedos D, Hewapathirana S, Kamatchinathan S, Kundu The PRIDE database resources in 2022: a hub for mass spectrometry-based proteomics evidences Nucleic Acids Res 50:D543–D552.

74. Mistry J, Chuguransky S, Williams L, Qureshi M, Salazar GA, Sonnhammer ELL, Tosatto SCE, Paladin L, Raj S, Richardson LJ, Finn RD, Bateman A. 2021. Pfam: The protein families database in 2021. Nucleic Acids Research 49:D412–D419.

75. Chen I-MA, Chu K, Palaniappan K, Ratner A, Huang J, Huntemann M, Hajek P, Ritter SJ, Webb C, Wu D, Varghese NJ, Reddy TBK, Mukherjee S, Ovchinnikova G, Nolan M, Seshadri R, Roux S, Visel A, Woyke T, Eloe-Fadrosh EA, Kyrpides NC, Ivanova NN. 2023. The IMG/M data management and analysis system v.7: content updates and new features. Nucleic Acids Research 51:D723–D732.

76. Madeira F, Pearce M, Tivey ARN, Basutkar P, Lee J, Edbali O, Madhusoodanan N, Kolesnikov A, Lopez R. 2022. Search and sequence analysis tools services from EMBL-EBI in 2022. Nucleic Acids Res 50:W276–W279.

77. Krogh A, Larsson B, von Heijne G, Sonnhammer EL. 2001. Predicting transmembrane protein topology with a hidden Markov model: application to complete genomes. J Mol Biol 305:567–580.

78. Sonnhammer EL, von Heijne G, Krogh A. 1998. A hidden Markov model for predicting transmembrane helices in protein sequences. Proc Int Conf Intell Syst Mol Biol 6:175–182.

79. Almagro Armenteros JJ, Tsirigos KD, Sønderby C K, Petersen TN, Winther O, Brunak S, von Heijne G, Nielsen H. 2019. SignalP 5.0 improves signal peptide predictions using deep neural networks. 4. Nat Biotechnol 37:420–423.

80. The UniProt Consortium. 2023. UniProt: the Universal Protein Knowledgebase in 2023. Nucleic Acids Research 51:D523–D531.

81. The PyMOL Molecular Graphics System, Version 2.0 Schrödinger, LLC.

